# Single-cell analysis of chromatin accessibility and gene expression in *Drosophila melanogaster* embryos following hypoxia treatment

**DOI:** 10.64898/2026.07.18.739350

**Authors:** Dan Zhou, Chenxu Zhu, Jin Xue, Caitlin Marsh, Tsering Stobdan, Bing Ren, Gabriel G. Haddad

## Abstract

Limited oxygen supply or hypoxia can impair fetal development and lead to developmental disorders, but the molecular mechanism underlying this phenomenon remains poorly understood. It is also well known that hypoxia results in transcriptomic alterations and epigenetic reprogramming. *Drosophila melanogaster* (fruit fly) has been used for decades as a powerful model to dissect the molecular mechanisms regulating development. To better understand the role of early hypoxic stress on development, we performed single-cell joint analysis of chromatin accessibility and transcriptome to characterize the influence of hypoxia on *Drosophila* embryonic development. We identified hypoxia-induced alterations in both gene expression and chromatin accessibility across 22 cell groups, especially in the genes regulating organogenesis and development of neuronal, tracheal, and muscular systems, including a reduction of germ cells, suggesting a long-lasting influence of hypoxic stress at an early embryonic stage on development and reproduction. In summary, this study demonstrates that early embryonic hypoxia induces cell type- and dose-dependent changes in chromatin accessibility and gene expression, leading to distinct developmental phenotypic responses, such as reduced number of germ cells under both 3% and 5% O₂. We further conclude that the *tramtrack* (*ttk*) gene is critical in germ cell development and reproduction in *Drosophila melanogaster*.

## Introduction

Developmental factors (such as maternal, placental, and fetal factors) often reduce oxygen supply to the fetus (i.e., fetal hypoxia), which may impair fetal development and increase the risk of perinatal and infant mortality as well as disorders at later developmental stages (1-6). Although fetal cells can tolerate hypoxia to a certain level, cellular responses to counter severe and prolonged hypoxia are often inadequate and this may lead to injury (7-10). In addition, it has been well recognized that the cellular response to hypoxia depends on the cell type and differentiation stage. Furthermore, fluctuation of oxygen levels may result in transcriptome alterations through epigenetic reprograming (11-13).

*Drosophila melanogaster* (fruit fly) has been used extensively as a powerful model to dissect the molecular basis of animal development. The rapid generation time, the excellent and numerous genetic and molecular tools have made the fly indispensable for developmental basic research. For example, the in-depth analysis of genes involved in embryonic development in flies provided a foundation for developmental biology (14). Importantly, due to evolutionary conservation, many of the mechanisms discovered in fruit flies turned out to be homologous to those regulating human development and disease. Indeed, a considerable amount of knowledge regarding human development, aging, tumor formation, alcohol intoxication, neurodegeneration as well as hypoxia tolerance or susceptibility were obtained from studies in *Drosophila melanogaster* (15-22).

Previous studies indicated that hypoxia has a major effect on embryonic growth and development. For example, hypoxia induces a delay or a reversible arrest of cell cycle in *Drosophila* embryos depending on the severity and duration of the hypoxic stress, hence affecting growth in embryos (23-26). Since hypoxia can have different effects on different types of cells, we used a high-throughput method for a joint analysis of open chromatin and transcriptome (i.e., Paired-seq (27, 28)) in single cells. We detected distinct hypoxia-induced alterations in both gene expression and chromatin accessibility, especially in the genes regulating organogenesis and development of the neuronal, tracheal, and muscular systems as well as a remarkable reduction of germ cells, suggesting a particular influence of severe hypoxic stress on brain, muscles and germ cells.

## Results

### Experimental design and data analysis

Embryos of the *Canton-S* fruit flies (wild-type *Drosophila melanogaster* line) were used in this experiment (29). The embryos were collected in a 2-hour time window between 12:00PM to 4:00PM and treated with either 3% O_2_ (an lethal O_2_ level that induces a delay or reversible arrest of cell cycle in *Drosophila* embryos (25)) or 5% O_2_ (an O_2_ level that dramatically reduces the survival of fruit flies (30)) or with normoxic room air (21% O_2_, as control) for 2 hours in room temperature. The embryos were collected and snap-frozen with liquid nitrogen following dechorionation. Next, the single nuclei suspension was prepared from frozen embryos using Dounce homogenizer. To generate Paired-seq library, 2.4 millions of permeabilized nuclei were subjected to the Paired-seq procedure as previously described (28) (Fig. 1A and 1B). To ensure reproducibility and sufficient numbers of nuclei recovered from the three conditions, we performed three Paired-Tag experiments from independently collected embryos. After sequencing, we removed low-coverage nuclei (< 300 transcripts and < 300 unique open chromatin reads per cell), and recovered 38,114 nuclei, with the median numbers of 1,207 UMI and 1,994 open chromatin reads per nuclei: corresponding to 9,404, 17,682 and 11,028 nuclei from the room air-treated (RA, control), 5% O_2_-treated (HYP5) and 3% O_2_-treated (HYP3) embryos, respectively. Next, we performed single-cell clustering with the Seurat software (31). Briefly, 500 variable genes across cells were selected with the Seurat software and dimensional reduction with principal component analysis (PCA, https://doi.org/10.1080/14786440109462720) were then performed, followed by clustering with the Louvain algorithm (https://iopscience.iop.org/article/10.1088/1742-5468/2008/10/P10008) and visualization with Uniform Manifold Approximation and Projection (UMAP, https://arxiv.org/abs/1802.03426). These nuclei were grouped into 22 unique clusters (Fig. 1C, 1D and Table S1) (31-35). Using specific signature gene markers, these clusters were annotated into 1 blastoderm cluster, 5 ectoderm clusters, 2 epidermis clusters, 1 gap cells cluster, 1 germ cell cluster, 1 hindgut proper primordium cluster, 6 muscle clusters and 4 neuronal clusters. Each cluster contains a different number of cells representing their proportion in the embryo. Among them, “Ectoderm-1” cluster is the largest cluster that represents 19.9% of the cells in RA embryos, 13.0% of the cells in HYP5 embryos, and 16.2% of the cells in HYP3 embryos. Whereas “Muscle-1” cluster is the smallest cluster that represents < 1% of cells under all 3 conditions (Table S1, Fig S1).

**Figure 1.**
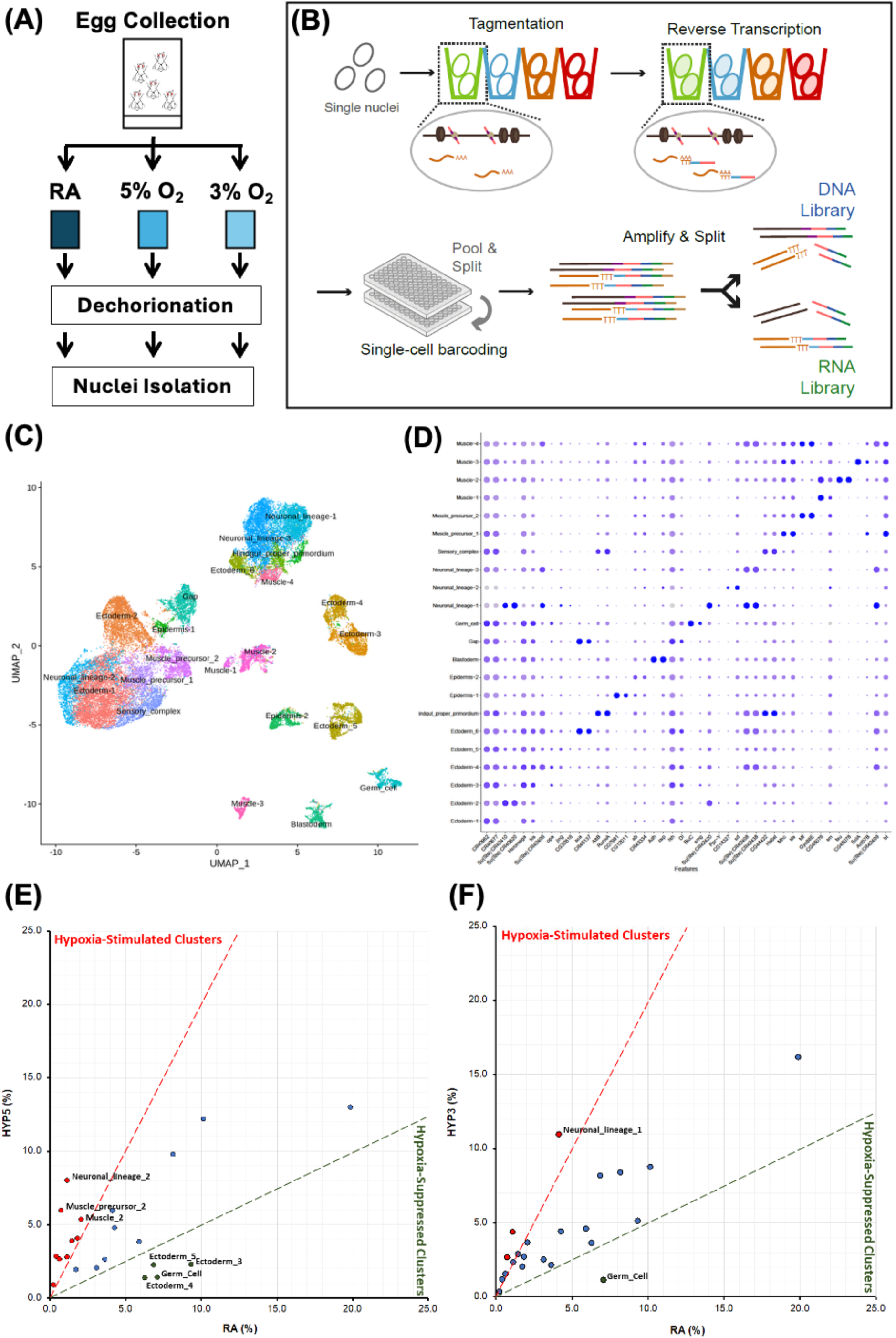
Experimental design and single-nucleus multi-omics analysis of Drosophila embryos under hypoxic stress. (A) Experimental workflow. Embryos of the Canton-S *Drosophila melanogaster* line were collected over a 2-hour time window, treated with room air (RA, 21% O₂), 5% O₂ (HYP5), or 3% O₂ (HYP3), dechorionated, snap-frozen, and processed for nuclei isolation. (B) Schematic of the Paired-seq workflow. Permeabilized nuclei were subjected to tagmentation and reverse transcription, followed by single-cell barcoding, amplification, and library preparation to generate paired DNA accessibility and RNA expression profiles. (C) UMAP visualization of 38,114 nuclei passing quality control, grouped into 22 distinct clusters based on gene expression signatures. (D) Dot plot showing a subset of signature marker genes used to annotate clusters into 1 blastoderm, 5 ectoderm, 2 epidermis, 1 gap cell, 1 germ cell, 1 hindgut primordium, 6 muscle, and 4 neuronal clusters. (E and F) Hypoxia-induced cell type-specific changes. Scatter plots comparing cluster proportions between RA and HYP5 (E) or RA and HYP3 (F). Clusters with >2-fold increase in proportion under hypoxia are identified as “hypoxia-stimulated clusters” (red) (including *Muscle-precursor-2*, *Neuronal-lineage-2*, *Muscle-4*, and *Muscle-3*), whereas clusters with >2-fold reduction in proportion under hypoxia are defined as “hypoxia-suppressed clusters” (green) (including *Germ cell*, *Ectoderm-3*, and *Ectoderm-4*). The *Germ cell* cluster showed the strongest suppression across both hypoxic conditions, whereas *Muscle-precursor-2*, *Neuronal-lineage-2*, *Muscle-3* and *Muscle-4* were strongly stimulated by hypoxia.

### Hypoxia treatment results in cell type-specific changes in early-stage embryos

A significant increase (or decrease) in cell number was detected in the HYP3 and HYP5 samples, suggesting that hypoxia induced a distinct, and cell type-specific response in *Drosophila* embryos. As shown in Figure 1E and 1F, the top clusters that had a suppression in their relative proportion (i.e., the hypoxia-suppressed clusters) are the “Germ-cell”, “Ectoderm-4” and “Ectoderm-3” clusters, and the top clusters that had a hypoxia-induced increase in their relative proportion (i.e., the hypoxia-stimulated clusters) include the “Muscle-precursor-2”, “Neuronal-lineage-2” and “Muscle-4” clusters. In general, clusters under HYP5 condition showed more changes than the clusters under HYP3 condition. The “Germ-cell” cluster was the top hypoxia-suppressed cluster whose relative proportion was decreased more than 2-fold under both HYP5 and HYP3 conditions. In contrast, the top clusters stimulated by hypoxia were “Muscle-precursor-2”, “Neuronal-lineage-2”, “Muscle-4” and “Muscle-3”, in which both hypoxic conditions induced a >2-fold increase in the relative proportion.

### Cell type-specific impact of hypoxia on the transcriptome and chromatin accessibility

Hypoxia treatment induced a wide range of alterations in transcriptome and chromatin accessibility among different clusters (Table S2). To determine the hypoxia-induced changes in transcriptome, a pseudo-bulk analysis strategy was applied following snRNA-seq to determine differentially expressed genes (DEGs) by comparing gene expression among the same cell clusters between hypoxia treatment and room air control. We found that, in the HYP3 embryos, the top 3 significant expressional changes were detected in the “Ectoderm-1” (total 1645 DEGs), “Neuronal-lineage-1” (total 1560 DEGs) and “Ectoderm-4” (total 1551 DEGs) clusters, but many fewer changes were detected in the “Muscle-precusor-1” (total 40 DEGs), “Muscle-4” (total 35 DEGs) and “Muscle-1” (total 22 DEGs) clusters. Interestingly, in the HYP5 embryos, the top significant expressional changes were detected in the “Neuronal-lineage-3” (total 2568 DEGs), “Ectoderm-1” (total 2267 DEGs) and “Ectoderm-2” (total 1841 DEGs) clusters.

To determine the changes in chromatin accessibility, we performed snATAC-seq and identified 53,000 – 65,000 peaks in the clusters with MACS2 software (36). Fifty percent in the accessibility peaks was differentially distributed in the genome in a condition-dependent manner following hypoxia treatment (Figure 2, Table S3). Noteworthy was that most of the regions that had a decrease in accessibility with hypoxia were the same in HYP5 and HYP3 groups. In contrast, the genomic loci with increased accessibility were more specific for the strength of hypoxic treatment (Fig. 2).

**Figure 2.**
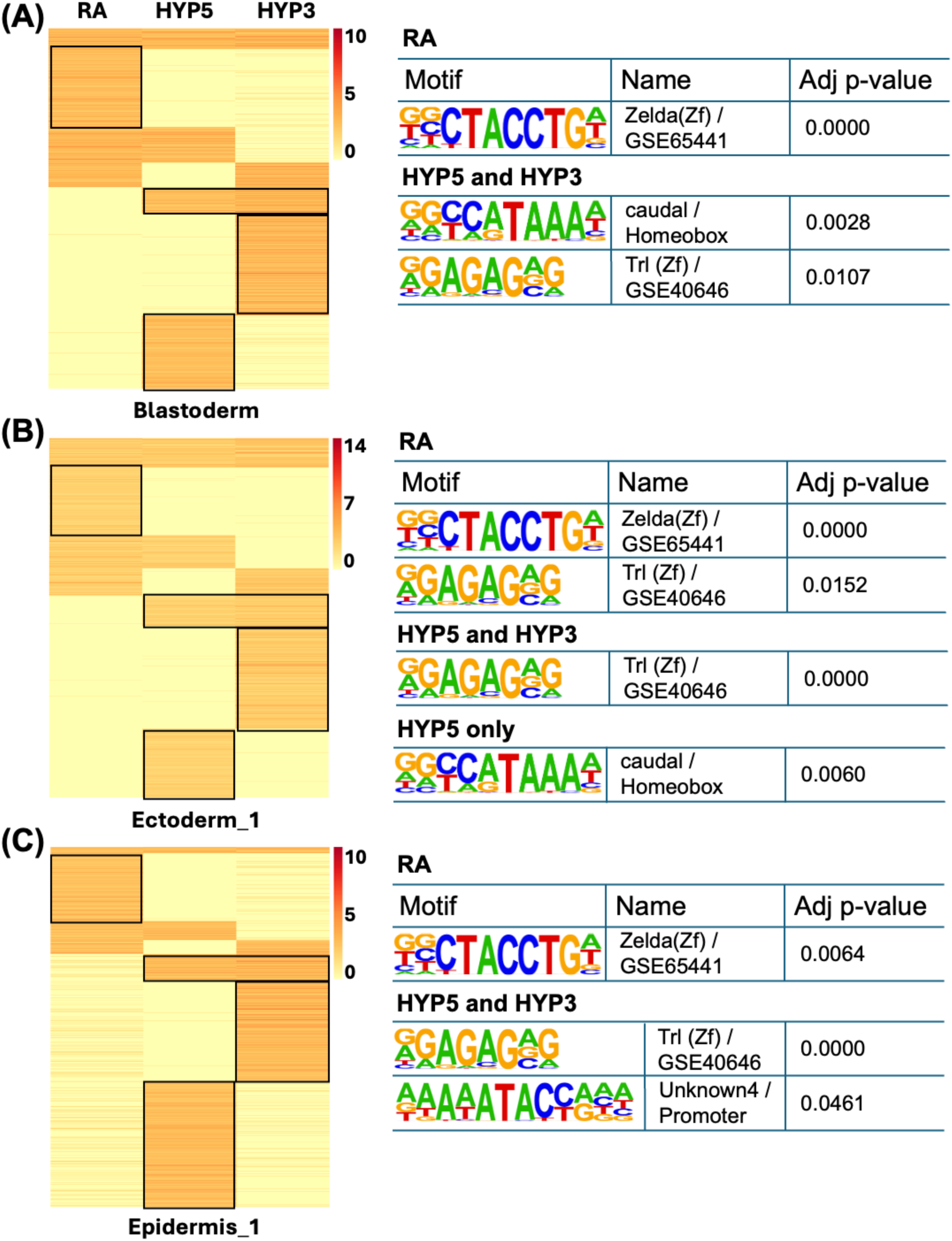
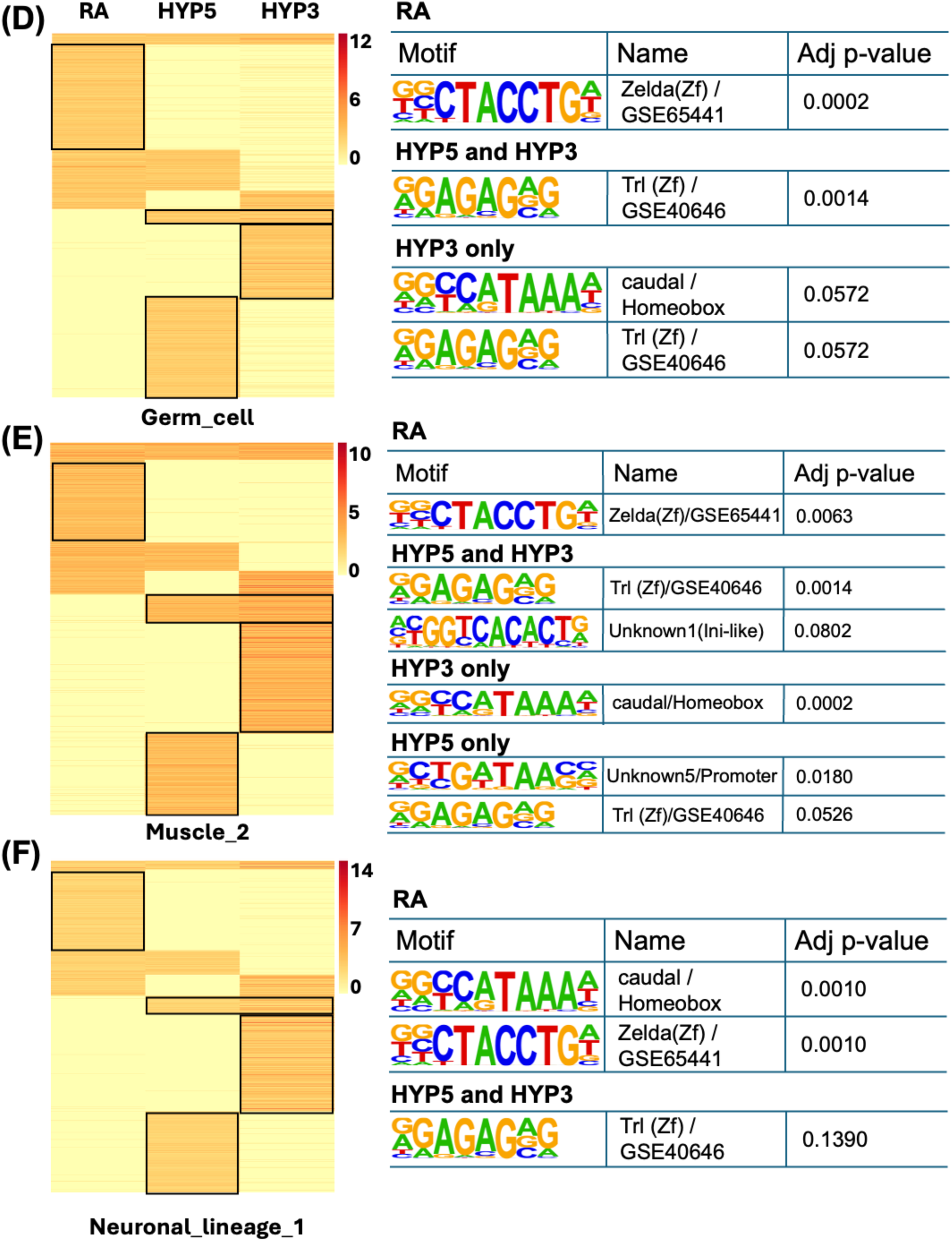
Hypoxia alters chromatin accessibility and transcription factor binding motif usage in specific clusters. Representative examples of hypoxia-induced changes in chromatin accessibility in the genome and motif enrichment across selected clusters. Heatmaps (left) show differential accessibility peaks identified by snATAC-seq under room air (RA), 5% O₂ (HYP5), and 3% O₂ (HYP3) conditions. Motif enrichment analysis (right) highlights transcription factor binding motifs associated with altered chromatin accessibility in each condition. (A) *Blastoderm* cluster (total 60725 peaks), (B) *Ectoderm-1* cluster (total 60588 peaks), (C) *Epidermis_1* cluster (total 53794 peaks), (D) *Germ cell* cluster (total 57292 peaks), (E) *Muscle_2* cluster (total 65543 peaks) and (F) *Neuronal_lineage_1* cluster (total 64507 peaks). Across these clusters, hypoxia treatment caused widespread reorganization of accessible chromatin regions, with many loci showing reduced accessibility common to both HYP5 and HYP3. Enriched motifs included those related to zygotic genome activation (e.g., Zelda) and anterior/posterior patterning (e.g., caudal). Notably, Trl and caudal motifs exhibited condition-specific relocation, with previously accessible regions in RA becoming less accessible and new loci opening under hypoxia, suggesting a strong influence of hypoxia treatment on early embryonic development.

A motif enrichment analysis was performed to determine the impact of alterations in chromatin accessibility on the binding elements of transcription regulators. We found that the most affected transcription regulator binding motifs are those related to chromatin modifications (e.g., Trithorax-like (Trl, GSE40646) and DNA replication-related element factor (Dref)), zygotic genome activation (e.g., Zelda (zld, GSE65441)) or anterior/posterior patterning formation (e.g., caudal (cad)) (Table S4). Among them, hypoxia treatment led to a shift of physical locations of Trl and cad binding motif on chromatin, where the opening locations under room air condition were reduced, and new locations became more accessible under hypoxia (Fig. 2 and Table S4). Furthermore, hypoxia treatment induced a reduction of the accessibility on the chromatin regions containing the initiator (INR) binding motif in the cells of 6 clusters (i.e., the clusters of Blastoderm, Ectoderm 3 to 5, Muscle_2 and Neuronal_lineage_3). In contrast, cells in 11 clusters of the HYP5 group (including the Ectoderm_5, Muscle_2 and Neuronal_lineage_3 clusters mentioned above) and the Ectoderm_3 cluster of the HYP3 group showed an increased accessibility of INR motifs at other chromatin regions with hypoxia-induced opening (Table S4), suggesting that HYP5 embryos had a stronger transcriptomic response through transcriptional activation potentially through rearrangements of chromatin accessibility.

### Concordant changes in chromatin accessibility and gene expression in embryonic development

We further investigated the influence of the changes in chromatin accessibility on the promoter regions and their correlation with the transcription level of the corresponding genes. Interestingly, hypoxia treatment altered the accessibility of hundreds of promoters in the genome of the hypoxia-suppressed clusters. In contrast, none or a few (<30) alterations were detected in the clusters that were stimulated by hypoxia (Table S5), reflecting different chromatin modifications in these two types of clusters/cells. Furthermore, although the promoter accessibility and the level of the corresponding transcripts exhibited unique heatmap patterns across clusters, the patterns correlated very well within each cluster (Fig. 3). And the correlation in the significantly altered promoter-DEG pairs (i.e., the genes with significantly altered promoter accessibility and transcription activity (|FC| > 1.5 and p < 0.05)) were very strong (>0.60 in the HYP3 and >0.85 in the HYP5 clusters, Table S6).

**Figure 3.**
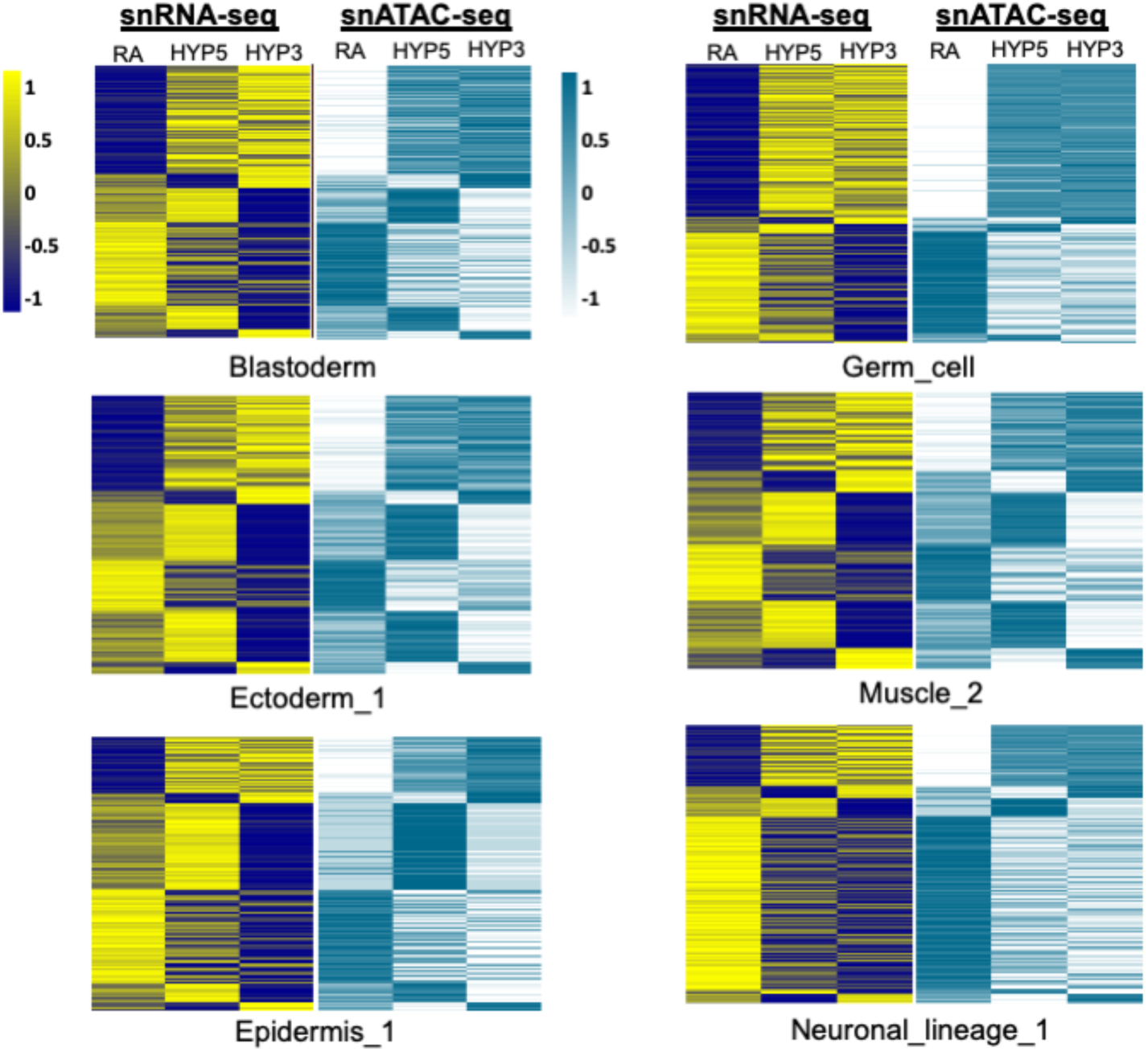
Synchronized alterations of chromatin accessibility and transcriptional activity and the impacts in biological processes following hypoxia treatment. Heatmaps show paired snRNA-seq (left of each pair) and snATAC-seq (right of each pair) signals for genes with significantly altered transcriptional activity (|FC| > 1.5 and p < 0.05) and promoter accessibility across multiple embryonic cell clusters, including Blastoderm (2627 pairs), Ectoderm_1 (3912 pairs), Epidermis_1 (763 pairs), Germ_cell (2046 pairs), Muscle_2 (1789 pairs), and Neuronal_lineage_1 (2577 pairs). Columns represent RA (normoxia), HYP5, and HYP3 hypoxia conditions. Despite distinct cluster-specific heatmap patterns, promoter accessibility and corresponding transcript levels showed strong within-cluster concordance. Correlations between significantly altered promoter–DEG pairs were high, exceeding 0.60 in HYP3 and 0.85 in HYP5, indicating a tight coupling between chromatin accessibility and transcriptional responses under hypoxic conditions.

We identified a total number of 266 Promoter-DEG pairs (increase: 135, decrease: 131) that were significantly altered in both HYP3 and HYP5 embryos and had significant alterations on both the accessibility of the promoters and the expression level of the transcripts (Table S6). Most of the changes were detected in the Ectoderm_1 to 6 and the Neuronal_lineage_3 clusters. It is interesting to note that none of these clusters exhibited a hypoxia-stimulated relative proportional change in either HYP3 or HYP5. According to GO (gene ontology) enrichment analysis, this set of 266 genes are involved in regulating cell differentiation, cell cycle and proliferation, which potentially influence the development of neuronal, tracheal, and muscle systems during *Drosophila* embryonic development (Table S6) (37-41).

### Hypoxia inhibits Germ cell development through epigenetic modification of ttk expression

Hypoxia induced a greater than 4-fold reduction of the Germ cell cluster in both HYP5 and HYP3 embryos (Fig. 1E and 1F), which begs the question as to whether this is a result of the hypoxia-induced changes in gene expression and the correlated alterations of the accessibility of their promoter regions. Indeed, collectively we detected 22 promoter-DEG pairs that showed concordant changes at both levels of hypoxia. Among these, 11 genes showed an up-regulated expression, and the other 11 genes had a down-regulated expression (Table S7). To evaluate functionally the effect of hypoxia on germ cells and reproduction, we used a germ line specific driver, nanos-Gal4, to express UAS-shRNA (or UAS-cDNA) transgenes targeting 18 candidate genes that have available UAS stocks in the germ cells. As shown in Fig. 4 and Table S8, knocking down (KD) *DopEcR* (*Dopamine/Ecdysteroid receptor*) or *rib* (*ribbon*) reduced the number of both F1 and F2 progenies. In contrast, *Nrt* (*Neurotactin*) or *ttk* (*tramtrack*) KD had no effect on F1 but increased the number of F2 progeny. Interestingly, an opposite effect was observed following ttk overexpression, in which the number of progenies in both F1 and F2 was reduced, which is consistent with the phenotypic reduction in progeny in both hypoxic levels (30, 42).

**Figure 4.**
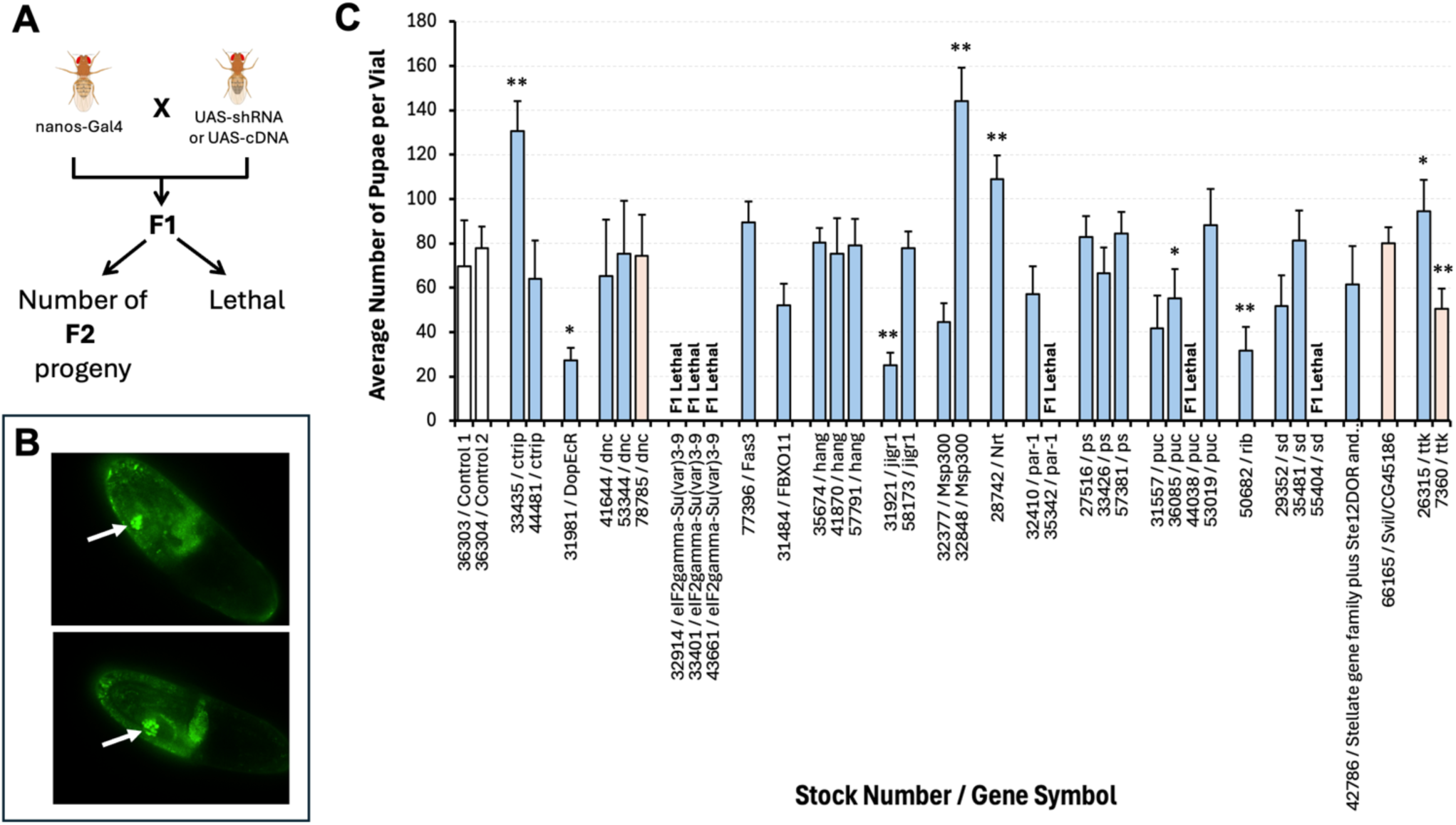
Germ line modification of DopEcR, Nrt, rib and ttk regulates reproduction power in *Drosophila*. (A) Schematic illustration of experimental design. (B) Evaluation of germline specificity of nanos-Gal4 driver. The nanos-Gal4 driver was crossed with UAS-GFP to label the cells with Gal4 expression (white arrow: germline cells). (C) The role of candidate genes in reproduction was tested by germline specific knocking down (blue bars) or overexpression (red bars). The F2 progeny was quantified using number of pupae and compared to control crosses (white bars). Knocking down *DopEcR* or *rib* had a significant reduction of F2 progeny; in contrast, knocking down *Nrt* increased the number of F2 progeny. An opposite effect was observed between *ttk* KD and OE, in which KD of *ttk* increased but OE of *ttk* decreased the number of F2 progeny, demonstrating the role of *ttk* gene in embryonic germ line development in *Drosophila melanogaster*. The statistical significance was determined with unpaired t-test (* p<0.01, ** p<0.001).

## Discussion

Mounting evidence has suggested that hypoxia has a major impact on embryonic development (11, 43, 44). In the current study, we found that acute hypoxia of two hours can induce cell-type specific responses in Drosophila embryos. As shown in Fig. 1, hypoxia treatment changed the relative cellular proportion in the embryo. For example, a remarkable reduction of cell proportion was observed in germ cells and 3 clusters of ectodermic cells. In contrast, an increase in relative cell proportion was found in the clusters of muscle cells and cells within different neuronal lineages, demonstrating that hypoxia treatment may lead to both arrest (or delay) and acceleration on cell cycle or proliferation depending on cell type. The remarkable reduction of the number of germ cells suggested a strong impact of early embryonic hypoxia on adult fly reproduction. In addition to the difference in transcriptome, such differential response further reflected the distinct epigenetic configurations between clusters.

Paired-seq combines snATAC-seq and snRNA-seq, allowing us to simultaneously determine the hypoxia-induced alterations in both chromatin accessibility and transcriptome in the same single nucleus. It is a powerful tool for understanding epigenetic modifications as well as transcriptional activities in each cell at the same time. Indeed, we detected significant changes in chromatin accessibility in both HYP5 and HYP3 embryos across almost half of the detected peaks in each cluster. Although most of the hypoxia-induced decrease of accessibility was detected in common genomic regions, the regions with increased accessibility were largely different, suggesting that, depending on the strength of hypoxic stress, hypoxia may induce chromatin opening at distinct genomic regions, which, in turn, may lead to activation of different sets of genes (Fig. 2 and 3).

The alterations in chromatin configuration changed the accessibility of multiple binding motifs for different chromatin modifiers and transcription regulators. Such alterations provided details about mechanisms that are induced by hypoxia, such as mechanisms regulating chromatin remodeling and transcription memory. For example, hypoxia reduced the accessibility of the binding motifs for Trl, zld, cad, Dref, M1BP, E-box and INR in the genomic regions where were higher accessible under RA condition, and increased the accessibility of Trl, cad (in HYP3) and Trl, cad, INR, M1BP, Dref, E-box (in HYP5) in other genomic regions (Fig.2 and Table S4). Trl (trithorax-like), the GAGA transcription factor, regulates transcriptional activity of many genes through modification of chromatin structure at their regulatory elements (45-47). It is also a critical regulator for oogenesis (48, 49). The oxygen-dependent switch of its genomic accessibility suggested that the Trl chromatin modifier plays a critical role in regulation hypoxia-induced chromatin remodeling, hence, transcriptomic response. Furthermore, the initiator (INR) motif is the most common core promoter element in metazoans that regulates activation and pausing of RNA polymerase and transcription activities of the developmental control genes (50-54). It has been shown that, in early *Drosophila* embryo, Polymerase II pausing is most frequently found at developmental control genes, and a large portion of these genes remain highly paused throughout embryonic development independently of their expression (55). Such pausing on RNA polymerase II may facilitate rapid temporal and spatial changes in gene activity during development (56). Further, it has been shown that environmental stresses (such as heat shock and oxidative stress) may switch RNA polymerase II from pausing to activation (57-59). For example, in mammalian cells, heat shock stress may form a “transcriptional memory” through promoter-proximal pause-release and pre-mRNA-processing (59). The current finding of increased accessibility of the INR elements in many more clusters treated with the milder hypoxia than those treated with the severe hypoxia (i.e., 11 clusters in the milder hypoxic HYP5 group versus 1 cluster in the severe hypoxic HYP3 group) suggested an oxygen-dependent and strength-sensitive epigenetic mechanism through differential modification of distinct genomic regions.

The spatiotemporal gene expression patterns of multi-cellular organisms are driven in large part by the cis-regulatory elements (CREs) in the genome (60). The Paired-seq approach allows us to better understand the cell-type specific gene regulatory programs by measuring the transcriptome and states of the candidate CREs within the same cells simultaneously. Using this tool, we found that the hypoxia-induced alterations in chromatin accessibility and transcription activity are well correlated. As shown in Fig. 3, the levels of accessibility of the CREs/promoter regions and the transcriptional activity of their putative downstream genes matched very well with an almost identical pattern. Since snRNA-seq measures the level of transcripts in the nucleus, it may directly reflect the “real-time” state of transcription activity of genes. Most likely, our data demonstrate the dynamic changes rather than the steady state homeostasis of the transcriptome following hypoxia treatment.

It is interesting to note that hypoxia-induced changes in the promoter-gene pairs were unique and specific to the treatment strength (i.e., 3% or 5% O_2_). This set of results further demonstrated a strength-dependent modification on different chromatin regions, hence, the transcriptional activity of a distinct set of genes. These significantly altered genes regulate a wide spectrum of biological functions, including cell cycle, differentiation and proliferation as well as chromatin organization, which affect organogenesis and development of neuronal, tracheal and muscular systems (Table S6).

Our results demonstrated a reduced number of embryonic germ cells along with significant changed promoter-DEG pairs in the same cluster under both HYP5 and HYP3 conditions, suggesting an effect of hypoxic stress on reproduction in the adult. Among the 18 genes that were tested, 4 genes (*DopEcR*, *Nrt*, *rib* and *ttk*) showed significant impacts on F1 and F2 progenies, especially the ttk gene that showed an opposite effect between KD and OE. The *ttk* gene encodes 2 transcription repressors that are required for normal embryogenesis and imaginal disc development (61-64). It has been shown that loss of *ttk* function transforms support cells to neurons, but *ttk* overexpression results in the reverse transformation in *Drosophila* PNS (65) and blocks glial development in *Drosophila* CNS (66). Much like the effect of ttk KD on early oogenesis (67), our results in hypoxia demonstrated that the level of ttk expression has a significant effect on reproduction (i.e., the up regulation of *ttk* in the germ cell cluster plays a key role in the reduction of germ cells).

In summary, the current study analyzed early embryonic hypoxia exposure-induced alterations on both chromatin accessibility and gene transcription at single nucleus resolution. Our results demonstrated cell type-specific and hypoxic strength-dependent alterations on epigenome and transcriptome. We also observed distinct developmental responses between different cell types represented by the relative proportion of each cell cluster in the embryos, demonstrating a context-dependent stimulative (or suppressive) effect of hypoxia on cell differentiation and proliferation in different type of cells. In addition, a reduction of germ cell population was detected in the embryos following hypoxia treatment, and hypoxia-induced transcriptional changes of *ttk* gene regulates the reproduction power in *Drosophila melanogaster*.

## Materials and Methods

### Collection of embryos, hypoxia treatment and isolation of nucleus for sequencing

The *Drosophila melanogaster Canton-S* strain was used in the experiment. Embryo collection and treatment were performed at room temperature. To remove the out-of-stage eggs, the adult flies were first transferred to a new bottle prior embryo collection and allow to lay eggs for > 3 hours. Then, the flies were transferred to apple juice-agar plates and allowed to lay eggs for 2 hours. These eggs were randomly separated to 3 groups and transferred into hypoxic chambers fed with 3% or 5% O_2_ (balanced with N_2_) or normoxic chamber fed with room air (as control) for 2 hr. After treatment, the embryos were dechorionated for 30 seconds in 50% solution of bleach for exactly two minutes and extensively washed with deionized water and rinsed with 0.05M Tris/HCl buffer (0.05M Tris/HCl; 25mM KCl; 350mM Sucrose; pH 7.6). Excess liquid was removed and embryos were snap frozen on dry-ice and store at - 80°C for nucleus isolation and sequencing.

### Paired-seq procedure

To prepare single-cell suspensions for Paired-seq assay, the frozen embryos were suspended in Douncing Buffer (0.25 M sucrose (Sigma, S7903), 25 mM KCl (Sigma, P9333), 5 mM MgCl_2_ (Sigma, 63069), 10 mM Tris-HCl pH 7.4 (Sigma, T4661), 1 mM DTT (Sigma, D9779), 1x Protease Inhibitor (Roche, 05056489001), 0.5 U/µL RNase OUT (Invitrogen, 10777-019), and 0.5 U/µL SUPERase Inhibitor (Invitrogen, AM2694), 0.1% Triton-X100 (Sigma T9284)) and dounced with douncer (KIMBLE, 885302) for 10-15 times, followed by filtering with 30 µm Cell-Tric (Sysmex). Cells were then spun-down for 10 min, at 1,000 g 4°C, washed with Douncing Buffer once and spun-down again. The cell pellets were then resuspended in Nuclei Permeabilization Buffer (10 mM Tris-HCl pH 7.4, 10 mM NaCl (Sigma, S7653), 3 mM MgCl_2_, 1× Protease Inhibitor, 0.5 U/µL RNase OUT, and 0.5 U/µl SUPERase Inhibitor, and 0.1% IGEPAL CA630 (Sigma, I8896)) and centrifuged for 10 min at 1,000g 4°C.

Paired-seq procedures were performed based on the previously described method (28) with minor improvements. Briefly, 2.4 million permeabilized nuclei were aliquoted into 12 tubes and spun down at 1,000g, 4 °C for 10 min. Nuclei were then resuspended in 18 ul of 1.1X Tagmentation Buffer (36.7 mM Tris-Ac, pH 7.8, 12.1 mM MgAc, 73.3 mM KAc, 17.8% DMF (Sigma, DX1730-6), 82.5 µM PitStop2 (Sigma, SML1169)). Barcoded Tn5 (1 µL, 0.5mg/mL) was then added, and the tagmentation reaction was carried out at 37 °C, 550 r.p.m. for 30 min in a ThermoMixer (Eppendorf). Reactions were quenched by adding 5 µL of 40 mM EDTA and nuclei were spun down again at 1,000g, 4 °C for 10 min and proceeded to reverse transcription as previously described (28).

### Paired-seq data analysis

1. Data preprocessing: Preprocessing and reads cleaning was performed with custom scripts available from GitHub (https://github.com/cxzhu/Paired-Tag). Cleaned reads were mapped to Drosophila reference genome Dm6 with STAR (v2.6.0a) and PCR duplicated reads were removed based on the mapping position, cellular barcode, and UMI. Low-coverage nuclei were removed from further analysis (< 500 transcripts and < 500 unique DNA reads).

2. Cell clustering: RNA alignment files were converted to cell-to-gene matrices with cells as columns and genes as rows. Cells with less than 300 UMIs and 300 open chromatin fragments detected were removed from clustering. Cell-to-gene counts were normalized, and variable genes were selected for dimensional reduction with principal component analysis (PCA) with the Seurat package. Batch effects were corrected with the harmony package, and cells were clustered with the Louvain algorithm. Cell cluster identities were annotated by FlyMine BDGP enrichment analysis of highly expressed genes (68).

3. Differential gene expression analysis: Read counts from all cells for each technical replicate of given clusters were summed were summarized, and differentially expressed genes (DEGs) were identified with edgeR software. To analyze the relationships between gene expression and promoter chromatin accessibility, DNA reads were aggregated according to RNA-clustering-based cell annotation and read per million total reads (RPM) were calculated for each gene promoter (-1.5kb to +500bp). To identify correlated gene-promoter pairs, DEGs with the same trends for both gene expression and promoter chromatin accessibility levels were selected.

4. Peak calling and motif analysis: DNA reads for each condition and cell type were aggregated based on RNA-clustering-based cell annotations. Peak calling was performed with the macs2 software with default parameters and overlapping peaks (peaks with at least one bp overlapped) were merged and extended to 1kb from merged peak centers. To group peaks into different modules, k means clustering was performed for reads densities in each peak across the three conditions in each cell group, and k = 7 was selected based on the reads densities relationships across different conditions. Motif analysis for each module was performed with Homer software with default parameters.

5. Gene Ontology Term Enrichment Analysis: The PANTHER Classification system was used for GO term enrichment analysis (69). All genes detected in the snRNA-seq were used as the background reference. For the cluster-specific GO term enrichments following hypoxia treatment, the list of genes identified as significant DEGs in a cluster (|FC| > 2 and p-value < 0.05) was applied.

### Germ-line specific silencing or overexpression of candidate genes

For germ-line specific silencing (or overexpression), the nanos-Gal4-VP16 driver (Bloomington Drosophila Stock Center #4937) was used to express shRNAs targeting candidate genes (or the cDNA transgenes of candidate genes), the nanos-Gal4-VP16/UAS-shRNA (or UAS-cDNA) flies were then screened for F1 survival and reproductivity. The survival rate of F1 was scored by number of pupae. To test the reproductivity of the F1 progeny, a single (nanos-Gal4/UAS-shRNA (or UAS-cDNA)) female was transferred into a new culture vial with 2-3 males of the same cross (10 vials for each candidate KD or OE) and let them lay eggs for 5 days. The number of pupae formed in each vial was scored and compared to the controls (70, 71).

## Data availability

The sequencing data obtained in this study have been deposited to the NCBI Gene Expression Omnibus (GEO) (http://www.ncbi.nlm.nih.gov/geo/) under accession number GSE330719 (*Reviewer access token: yxidcuaonvmxfqf*) and are publicly available as of the date of publication.

## Acknowledgements

This work was supported by NIH grant R21 NS120023 and R01 HL157445 to GGH.

